# Assessment of technical and clinical utility of a bead-based flow cytometry platform for multiparametric phenotyping of CNS-derived extracellular vesicles

**DOI:** 10.1101/2023.07.14.549082

**Authors:** Alexandra Brahmer, Carsten Geiß, Andriana Lygeraki, Elmo Neuberger, Theophilos Tzaridis, Tinh Thi Nguyen, Felix Luessi, Anne Régnier-Vigouroux, Gunther Hartmann, Perikles Simon, Kristina Endres, Stefan Bittner, Katrin S Reiners, Eva-Maria Krämer-Albers

## Abstract

Extracellular vesicles (EVs) derived from the CNS are potential liquid-biopsy markers for early detection and monitoring of neurodegenerative diseases and brain tumors. This study assessed the performance of a bead-based flow cytometry assay (EV Neuro) for multiparametric detection of CNS-derived EVs and identification of disease-specific markers. Different sample materials and EV isolation methods were compared. Glioblastoma- and primary human astrocyte-derived EVs exhibited distinct EV profiles, with signal intensities increasing with higher EV input. Analysis of serum or plasma from glioblastoma, multiple sclerosis, Alzheimer’s Disease patients and healthy controls showed varying marker signal intensities. Notably, data normalization improved marker identification. Specific EV populations, such as CD36^+^EVs in glioblastoma and GALC^+^EVs in multiple sclerosis, were significantly elevated in disease compared to controls. Clustering analysis techniques effectively differentiated glioblastoma patients from controls. A potential correlation between CD107a^+^EVs and neurofilament levels in the blood was identified in multiple sclerosis patients. Together, the semi-quantitative EV Neuro assay demonstrated its utility for EV profiling in complex samples. However, reliable statistical results in biomarker studies require large sample cohorts and high effect sizes. Nonetheless, this exploratory trial confirmed the feasibility of discovering EV-associated biomarkers and monitoring circulating EV profiles in CNS diseases using the EV Neuro assay.

## Introduction

Extracellular vesicles (EVs) recently gained considerable interest as minimally invasive diagnostic and prognostic biomarkers in a variety of diseases including central nervous system (CNS) pathologies (1–3). Subtypes of EVs sized between 50 nm and 1 µm are released by cells through different pathways including plasma membrane shed ectosomes, endosomal-derived exosomes, and blebbing apoptotic bodies. EVs function in cell-cell communication and, when released into the extracellular space, carry molecular cargo specific for the cell-type of origin and its cellular state (4). Recent evidence suggests that brain-derived EVs originating from neurons and glia can cross the blood-brain barrier (BBB) and appear among circulating EVs, where they can serve as biomarkers of neurodegenerative conditions or brain cancer (5–7). Neurodegeneration and brain tumors such as glioblastoma (GB), the most common malignant brain tumor (8), are commonly associated with neuroinflammatory conditions, known to promote BBB leakiness and EV shedding from brain microvascular endothelial cells. Therefore, brain-derived EVs possess high potential for minimal-invasive diagnosis and prognosis of diseases such as Alzheimer’s disease (AD), Parkinson’s disease, multiple sclerosis (MS) and GB, which currently are only detected when clinical symptoms arise and considerable damage has already occurred to the brain (9–13).

Circulating EVs in human plasma and serum represent a complex mixture of EVs derived from different tissues and cellular origins. The relative contribution of brain-derived EVs to the overall population of circulating EVs is unclear and expected to reflect a minor and dynamic fraction. Moreover, lipoprotein particles co-purify with and outnumber EVs by several orders of magnitude (14). Thus, brain-derived EVs must be revealed against a large background of potentially interfering particles. As a solution to this, immunocapturing of EVs via neuro-specific surface epitopes such as L1 cell adhesion molecule (L1CAM) or glutamate aspartate transporter (GLAST) has been used to enrich neuronal EVs or astrocyte-derived EVs, respectively (15–18). However, the specificity and efficiency of this strategy to depict brain-derived EVs is reported controversially, as these epitopes are also present on non-neural EVs or exist in soluble form (19,20). Therefore, multiparametric analysis of EVs can improve EV phenotyping and reveal a profile of EVs that may allow a comprehensive assessment of brain-derived EVs in complex samples.

Recently, a multiplex bead-based flow cytometry assay has been developed to perform a broad semi-quantitative profiling of EVs, covering up to 37 different EV surface epitopes (21,22). EVs are captured through specific antibodies on fluorescently bar-coded beads and detected in a second step using fluorescently conjugated antibodies recognizing genuine EV markers such as CD9, CD63, and/or CD81. The test can be performed with small sample input, does not require specific equipment and the readout can be performed with conventional flow cytometers. Moreover, an open-source software tool (multiplex analysis post-acquisition analysis software [MPAPASS]) has recently become available to enable high-throughput quality control and data analysis of multiplex EV data (23). The classic commercially available platform is designed to preferentially detect epitopes belonging to the cells associated with the human circulation (hematopoietic lineage, endothelial cells). A growing number of studies have used this platform to perform EV phenotyping and to reveal EV profiles and EV dynamics associated with different human conditions (21,24–26).

Here, we used a novel, prototype multiplex bead-based flow cytometry platform (MACSPlex EV Kit Neuro, human; short: EV Neuro) designed to detect brain-derived EVs that may be associated with CNS pathologies and evaluated the platform using different sample input materials. Marker performance and the semi-quantitative potential was first evaluated using EVs derived from cell culture supernatants including different human glioblastoma lines and primary human astrocytes. Furthermore, we performed an exploratory trial comparing EVs separated by size exclusion chromatography or immunoaffinity isolation from serum or plasma of healthy controls (HC) and patients with different CNS pathologies, including GB, MS, and AD. While detection and quantitative evaluation of individual CNS-specific markers remains challenging, the study shows that the multiparametric analysis may be useful to reveal EV marker profiles associated with distinct disease conditions.

## Materials and Methods

### Study design

In this study, a prototype multiplex bead-based EV Neuro platform (MACSPlex EV Kit Neuro, human, provided by Miltenyi Biotec, Bergisch Gladbach) was tested with different sample material and EV isolation protocols (Fig. 1). The prototype kit consisted of two panels and comprised 64 different markers, which included CNS epitopes typical for neurons and glia cells and more ubiquitous epitopes relevant to the CNS (such as integrins) next to the genuine EV markers CD9, CD63, and CD81. GB and astrocyte cell culture-derived EVs were isolated via differential ultracentrifugation. GB and HC serum-derived EVs were prepared using size-exclusion chromatography combined with UC and compared to healthy control serum-derived EVs. Moreover, blood plasma samples from MS patients, HC, and patients suffering from mild cognitive impairment (MCI), AD, depression (DEP), or a combination of the named, further referred to as MAD cohort, were prepared using immuno-affinity capture (CD63^+^EV or CD81^+^EVs). The isolated EVs were analyzed using the EV Neuro assay. Thus, the performance of the EV-Neuro kit was tested on different combinations of starting material and methods of EVs isolation.

**Figure 1:**
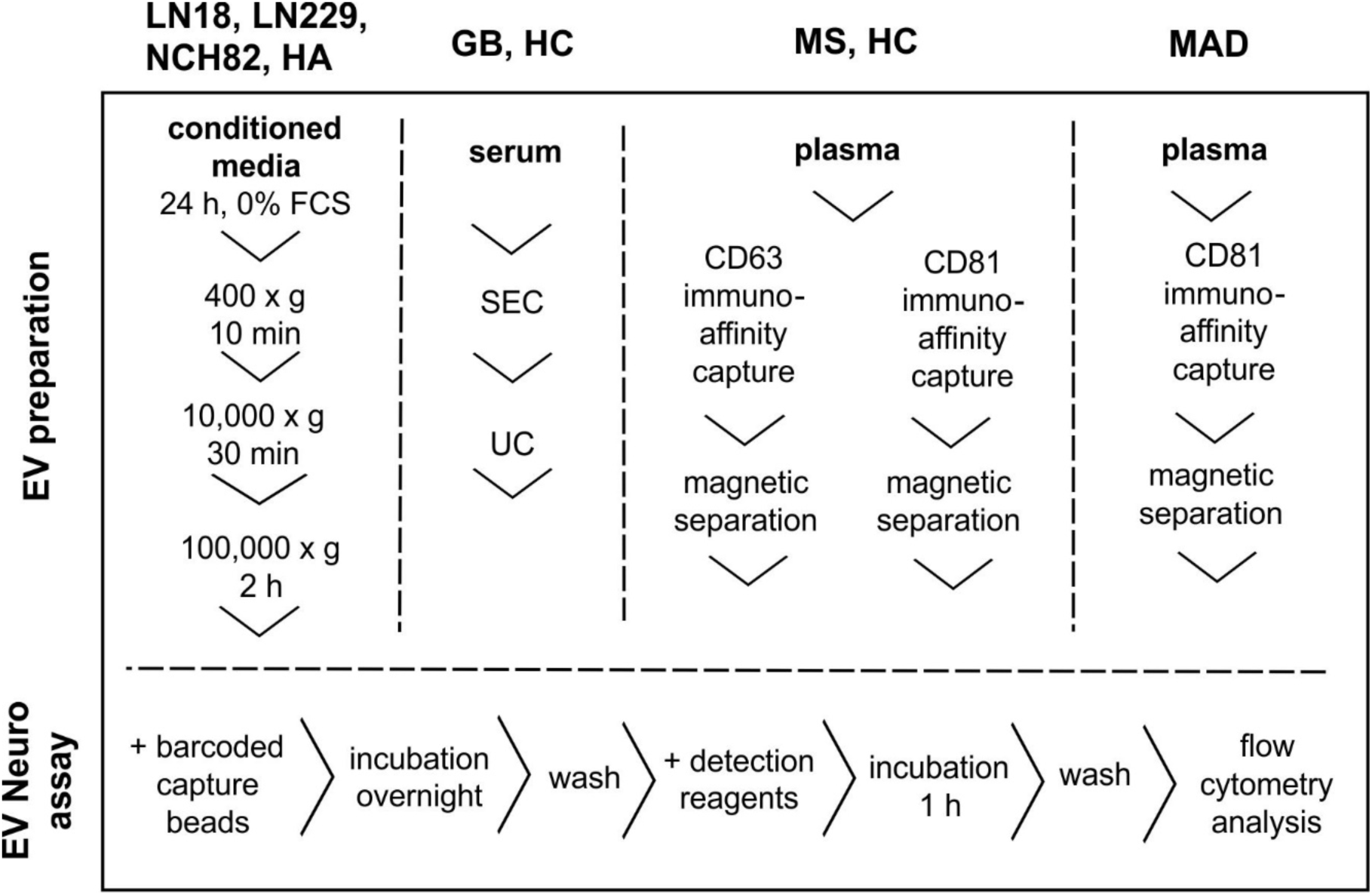
Graphic illustration of the different combinations of sample material and EV isolation protocols applied to the EV Neuro assay. For further details, see text. Abbreviations: HA: primary human astrocytes; GB: glioblastoma; HC: healthy control; MS: multiple sclerosis; MAD: mild cognitive impairment/ Alzheimer’s disease/depression; SEC: size exclusion chromatography; UC: ultracentrifugation.

### Participants and Ethics

9 GB patients [male (m): 7, female (f): 2, age: 46.4 (mean) ±14.8 (SD)] and 12 healthy persons [m: 7, f: 5, age: 52.6 ±13.4 y] were recruited at the Institute of Clinical Chemistry and Clinical Pharmacology, University Hospital Bonn. 11 MS (m: 2, f: 9, age: 44.0 ±11.0 y) patients and 5 healthy persons (m: 1, f: 4, age: 44.0 ±13.0 y) were recruited at the Department of Neurology, University Medical Center Mainz. 9 MAD patients (m: 5, f: 4, age: 68.8 ±8.6 y) were recruited at the Department of Psychiatry and Psychotherapy, University Medical Center Mainz. The samples of two MS patients and one MAD patient had to be excluded because of technical reasons (clogging of columns during immuno-isolation of EVs due to unknown reasons).

Informed written consent was obtained from all donors. The experimental procedures were approved by the Human Ethics Committee Rhineland-Palatinate and the local ethics committee of the University of Bonn, respectively, and adhere to the standards of the Declaration of Helsinki of the World Medical Association.

### Blood sample collection

For plasma preparation, venous blood was drawn using EDTA blood collection tubes and processed within 15 min. Platelet-free plasma was prepared by two rounds of centrifugation at 2,500 × g and room temperature (RT) for 15 min and stored at −80 °C until further processing.

For serum preparation, blood was collected in 9 ml S-monovettes (Sarstedt, Nuembrecht, Germany) and samples were processed as previously described (9). In short, samples were rested in an upright position for 30 min to allow coagulation and then centrifuged at 2,000 × g at RT for 15 min. Serum was transferred into fresh 15 ml tubes and centrifuged at 3,200 × g and RT for 20 min to remove platelets and stored at −80°C until further processing.

### EV separation from serum samples

EVs were isolated from 500 µl serum samples following the previously described method (9). Briefly, the serum was pre-cleared by centrifugation at 10,000 × g for 45 min at 4 °C. SEC was then performed using sepharose-based qEV columns (iZON Science, Christchurch, New Zealand) and the EVs were eluted with Hank’s balanced salt solution (HBSS). Fractions 8-10, each measuring 500 µl, were collected, pooled, and supplemented with a protease inhibitor (Roche, Basel, Switzerland). Subsequently, the samples were concentrated to a final volume of 240 µl through ultracentrifugation at 42,300 rpm using a Beckman TLA-55 rotor (RCF_avg_: 80,160; RCF_max_: 110,220; k-factor: 110 and the Optima MAX-XP ultracentrifuge (both Beckman Coulter, Brea, CA, USA) and equal sample volumes were immediately used for the EV Neuro panel A and B (115±25 µl, each).

### EV separation from plasma samples by immuno-affinity capture

1 ml plasma was thawed at RT and diluted in 1 ml PBS (MAD samples) or 3 ml PBS (MS and HC samples). 100 µl of CD81- (for all samples) or CD63- (for MS and HC samples) isolation beads (Exosome Isolation Kit, human Miltenyi Biotec) were added to 2 ml of diluted sample, which was then incubated at RT for 1 h under constant shaking. CD63^+^EVs and CD81^+^EVs were magnetically captured and eluted in 120 µl isolation buffer according to the manufacturer’s information using µ Columns and a µMACS™ Separator. The column-flowthrough of the sample was then loaded on a second µ Column (Miltenyi Biotec) and residual CD63^+^EVs and CD81^+^EVs, respectively, were magnetically captured and eluted in 120 µl isolation buffer. Consecutive eluates were pooled, split again in 120 µl each and immediately used in the EV Neuro assay. This resulted in input volume of plasma per EV Neuro panel A or B (see below) of 250 µl for MS as well as HC samples and 500 µl for MAD samples.

### EV separation from cell culture by differential ultracentrifugation (dUC)

LN18, LN229, and NCH82 glioblastoma cell lines were grown in cDMEM containing 10% FCS, 2 mM Glutamine, and 50 µg/ml Gentamycin. Cell lines were authenticated using Multiplex Cell Authentication by Multiplexion GmbH (Heidelberg, Germany). The SNP profiles matched known profiles or were unique. Primary human astrocytes (ScienCell, Carlsbad, CA, USA) were grown in AM medium containing 2% FCS, 1% astrocyte growth supplement (ScienCell), and 50 µg/ml Gentamycin. Before media collection, all cells were cultured in T75 flasks until they reached 70-80% confluency. Cells were washed with cDMEM without FCS and then cultured for 24 h in cDMEM without FCS until media collection. The cell culture supernatant was collected and pre-cleared from cells and larger particles by centrifugation at 400 × g at 4 °C for 10 min (Eppendorf 5810R) and 10,000 × g at 4 °C for 30 min (Eppendorf 5910R). Pre-cleared supernatants were subjected to ultracentrifugation at 29,000 rpm using a Beckman SW40 Ti rotor (RCF_avg_: 106,154; RCF_max_: 149,576; k-factor: 260) at 4 °C for 2 h. The dUC pellets were stored at −20 °C until further processing. The cells were trypsinized, washed in PBS, and cell lysates were prepared using RIPA buffer and stored at −20°C until further processing. Experiments were performed once for LN18 and LN229, twice for HA and three times for NCH82.

### Nanoparticle tracking analysis (NTA)

EVs isolated from cell culture supernatant were analyzed at 23°C using the Nanosight LM10 system (camera model Hamamatsu C11440-50B/ A11893-02; 532 nm laser). Nanosight 2.3 software (Malvern, Herrenberg, Germany) settings were: camera control in standard mode (camera level 14), particle detection in standard mode (detection threshold 8 and minimum expected particle size auto), and script control (Repeatstart, Syringeload 500, Delay 5, Syringestop, Delay 15, Capture 30 and Repeat 4). Five videos of 30 s were recorded per sample, particles were tracked (batch process), and average values were calculated. EV samples were diluted in particle-free PBS and measured in a range of 5–10 × 10^8^ particles/ml.

### Western blotting

EV pellets derived from 5.75 ml of conditioned media were resuspended in 1:1 diluted 4x sample buffer (200 mM Tris-HCL, pH 6.8; 10% SDS; 0.4% bromophenol blue; 40% glycerol; 400mM DTT; non-reducing conditions for CD9 and CD63 antibodies) and 20 µg of cell lysates were mixed with 4x sample buffer. The samples were subjected to SDS-PAGE (10 or 12% polyacrylamide gels) and Western blotting using PVDF membranes. The membranes were blocked with 4 % milk powder, 0.1 % Tween in TBS and incubated with primary and HRP-coupled secondary antibodies. Subsequently, proteins were detected utilizing chemiluminescence (SuperSignal™ West Pico PLUS Chemiluminescent Substrate, thermo scientific, Rockford, US) and X-ray films.

The following antibodies were used: CD9 (1:2000 dilution, clone #MM2/57, Merck Millipore, Darmstadt, Germany), CD63 (1:500 dilution, #CBL553, Merck Millipore, Darmstadt, Germany), CD81 (1:1000 dilution, #B-11, Santa Cruz, Heidelberg, Germany), Syntenin (1:3000 dilution, polyclonal, ab19903, Abcam, Cambridge, UK), Calnexin (1:4000 dilution, polyclonal, SPA-865, Stressgen, San Diego, US) and HRP-coupled secondary antibodies (Goat-anti-Mouse-HRP, 115-035-003, 1:10,000; Goat-anti-Rabbit-HRP, 111-035-003, 1:10,000; Dianova, Hamburg, Germany).

### Multiplexed bead-based flow cytometry assay

MACSPlex EV Kit Neuro, human, reagents were kindly provided by Miltenyi Biotec (Bergisch Gladbach). The marker composition of the prototype MACSPlex EV Kit Neuro, divided into two panels (A and B), can be found in table S1. Note that for GB serum EV analysis, the composition of the kit differed from the kit composition used for the other samples since some capture antibody beads (CD11b_2, CD146, NEFH, PVALB, SYP, TH, TUBB3, mIgG1_ctrl, REA_ctrl) were added to an updated prototype kit after measurement of GB serum samples. 120 µl of EV samples or pre-cleared cell culture supernatant were incubated with the panel A or B capture beads. The samples and a buffer control were processed according to the manufacturer’s instructions of the MACSPlex Exosome, human, Kit (Miltenyi Biotec). Briefly, after overnight incubation samples were washed and incubated for 1 h with a mix of the APC-labelled antibody-mix of anti-CD9, anti-CD63, and anti-CD81. Analysis for plasma- and cell line-derived EV samples was performed using the MACSQuant^®^ Analyzer 16 (Miltenyi Biotec) and the corresponding software MACSQuantify™ (Version 2.13.3). Data of serum-derived EV samples was acquired with the Attune NxT (LifeTechnologies, Darmstadt, Germany) and analyzed with FlowJo software, version 10 (BD Biosciences, San Jose, CA, USA). Signal intensity measured for each target was subtracted by the signal intensity for the respective target in the buffer control measurement. Negative values were set to zero. See workflow and gating strategy in Fig. S1.

### Measurement of neurofilament and GFAP levels in blood

Blood neurofilament (NFL) levels were measured in duplicates using the single molecule array HD-X analyzer (Quanterix, Boston, MA) and the NF-light Advantage Kit according to the manufacturer’s protocol.

### Data analysis

Raw data was assembled in Microsoft Excel 365 and further analyzed in R (version 4.1.2) using R Studio (version 2023.06.0). R packages tidyverse (version 2.0.0) and rstatix (version 0.7.2) were used for statistical analysis. Normal distribution of the data was tested with Shapiro-Wilk test. Unpaired samples t-test (normally distributed data) or Wilcoxon rank sum test (non-normally distributed data) were computed to identify statistically significant differences. ggplot2 (version 3.4.2) was used for data visualization. ComplexHeatmap package (version 2.10.0) was used for generation of heatmaps including cluster analysis using Pearson’s distance method. Rtsne package (version 0.16) was used for t-distributed stochastic neighbor embedding (tSNE) analysis. EnhancedVolcano package (version 1.12.0) was used for correlation analysis using Spearman’s rank correlation coefficient for non-normal distributed data. A p-value < .05 was considered significant. Parts of Supplementary Fig. 1 and the graphical abstract were drawn by using pictures from Servier Medical Art. Servier Medical Art by Servier is licensed under a Creative Commons Attribution 3.0 Unported License (https://creativecommons.org/licenses/by/3.0/).

### EV-TRACK

We have submitted all relevant data of our experiments to the EV-TRACK knowledgebase [EV-TRACK ID: EV230955 (27)].

## Results

### Qualitative assessment of GB cell line-derived EVs

To elaborate the general performance of the EV Neuro assay in a defined sample material, we analyzed EVs derived from three different astrocytic GB cell lines (LN18, LN229, and NCH82) and primary human astrocytes (HA) as healthy control cells. EVs were separated from cell culture supernatants using dUC and characterized via nanoparticle tracking analysis (NTA), showing the expected particle distributions with size peaks around 150 nm (Fig. 2a). As a quality control, we performed Western Blot analysis of representative EV-samples derived from the GB cells and HA, which confirmed the abundance of genuine EV markers CD9, CD63, CD81, and Syntenin (Fig. 2b). The non-EV marker Calnexin was found in NCH82- and LN18-EVs, though, not enriched in the EV samples. Next, we analyzed EVs isolated from equal volumes of cell culture supernatant of the different cell lines in the EV Neuro assay (Fig. 2c). Overall, the signal intensities obtained for the individual markers reflected NTA particle concentration, demonstrating that the assay can depict varying amounts of EVs and provide quantitative information.

**Figure 2:**
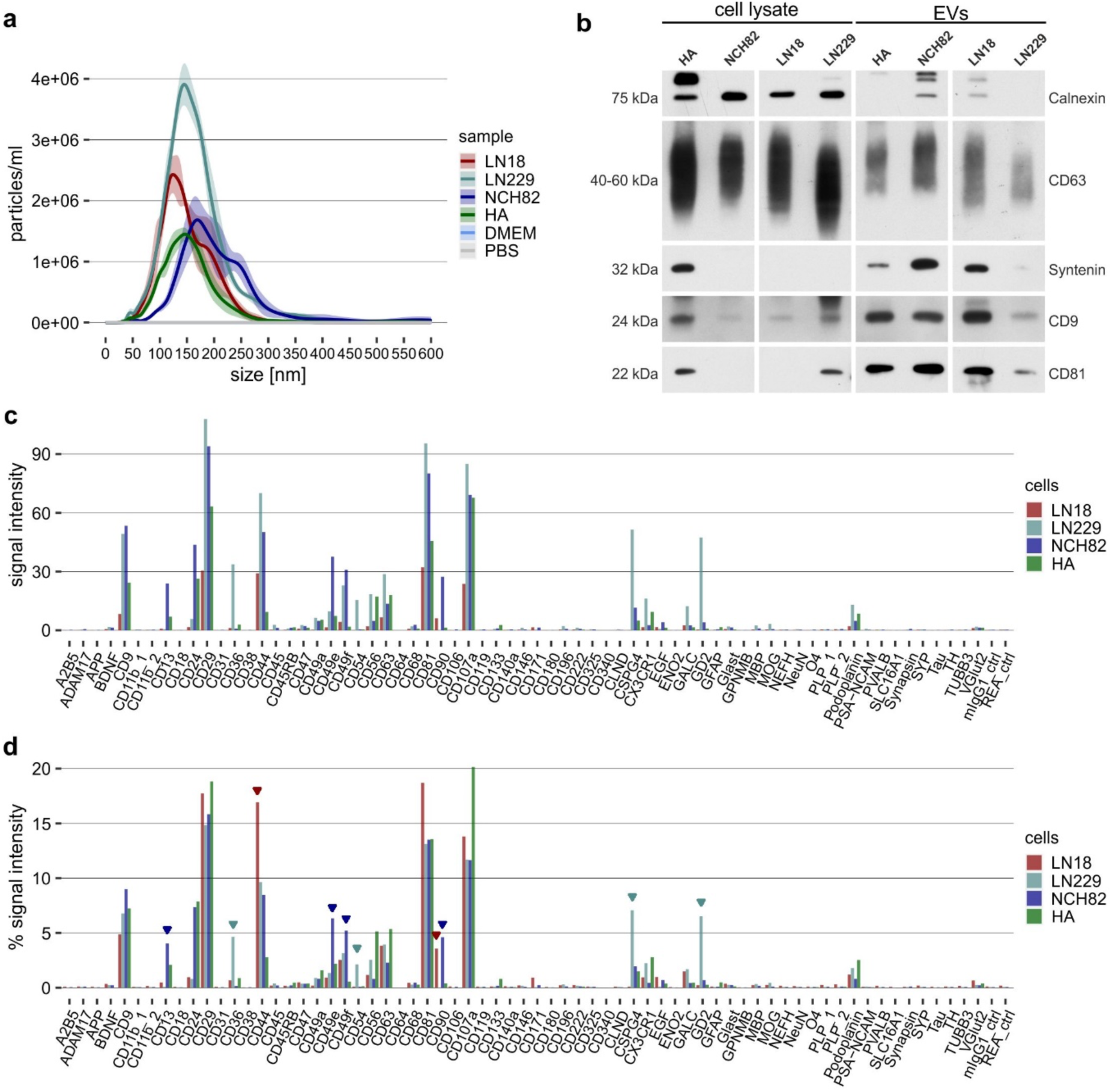
Characterization of EVs derived from glioblastoma cells and astrocytes. (a) Size distribution of EVs separated from cell culture supernatant of LN18, LN229, and NCH82 glioblastoma cell lines as well as from primary human astrocytes (HA). (b) Western blot analysis of CD9, CD63, CD81, Syntenin (EV markers) and Calnexin (non-EV marker) in EVs separated from cell culture supernatant of NCH82 glioblastoma cells and primary astrocytes. (c) EV Neuro median signal intensities for LN18, LN229, NCH82, and HA EVs as well as media control (DMEM w/o serum). (d) Relative signal intensities of values for cell derived EVs shown in (c). Relative signals are calculated by dividing the signal intensity of a target by the sum of intensities revealed from all markers (total intensity) and presented in percentages. Bars reflect representative marker profiles (n = 1-3 biological replicates). Arrow heads indicate markers that appear elevated in cell lines compared to primary astrocytes.

To better compare the EV profiles and depict markers that are specific for certain cell lines, we calculated relative signal intensities defined as signal of a target divided by the total of all signal intensities (Fig. 2d). Implementing this normalization strategy enables the visualization of markers that are equally represented within EVs of all cell lines (e.g., CD9, CD29, CD63, CD81) compared to markers that are distinct or even unique for a cell line. Accordingly, a relative increase of certain markers can be detected in the profile of cancer cell-derived EVs compared with normal primary astrocyte-EVs: CD44 and CD90 were elevated in LN18-EVs; CD36, CD54, CSPG4, and GD2 in LN229-EVs; CD13, CD49e, CD49f, and CD90 in NCH82-EVs. CSPG4, CD36, CD44 and GD2 were already shown to be involved in cancer cell migration and/or proliferation (28–31). Thus, EVs carrying these markers could indeed serve as biomarkers for glioblastoma.

However, markers such as the astrocytic Glutamate transporter GLAST or the intermediate filament glial fibrillary acidic protein (GFAP), which are expected to be present in GB or astrocyte-derived EVs were near to background or not detectable in EVs examined here. Background signal intensities for each capture bead population (CB) after incubation with detection antibody cocktail (DA) were always close to zero (signal intensity < 2) in several independent experiments (Fig. S2a), indicating absence of unspecific binding. Notably, the number of detected barcoded beads used for estimation of median signal intensities in flow cytometry measurement was strikingly low for some targets, which might hamper reproducibility of obtained data for these targets (Fig. S2b). As expected, all markers detected in the assay reflect surface epitopes, while cytoplasmic epitopes (e.g. NeuN, GFAP, MBP), which are contained within the EV lumen and thus not accessible for bead binding, never revealed signals.

Together, these findings highlight the EV Neuro assay’s ability to distinguish between different EV phenotypes by multiplexed marker analysis and to unveil surface markers that could qualify as biomarkers to be used for GB detection or characterization.

### Semi-quantitative comparison of GB-derived EVs

To further assess the semi-quantitative potential of the EV Neuro assay, we used EVs derived from NCH82 and LN18 at three different orders of magnitudes (Fig. 3). Input of native EVs in pre-cleared cell culture supernatant (120 µl, ∼3×10^7^ particles) did not reveal signals above background levels. After dUC enrichment of EVs, input of ∼3×10^8 and ∼1×10^9^ particles resulted in increasing signal intensities, though, the increases were not linear and differed between markers. A three-fold increase in particle input did not equally increase signal, which is most evident for markers with high signal intensity such as CD29 and CD81, indicating a potential saturation at higher marker concentrations. Notably, it was possible to detect further markers clearly above background when increasing the particle input (e.g., CD47, CD49 a, CD56, CSPG4, GD2). Still, GLAST was only detectable at low signal intensities, indicating either poor assay performance for this target or a very low amount of GLAST-carrying EVs in the samples. Importantly, relative signal intensities appeared to be largely independent of particle input (in particular those in the medium signal intensity range that probably perform in the linear detection range of the assay), confirming that normalization reveals an EV profile typical for the cell type of origin (Fig. S3). In conclusion, the EV Neuro assay can be used to screen defined EV populations and is able to depict differences in a semi-quantitative manner. Moreover, increasing input EV concentration can result in further marker detection. Multiparametric phenotyping reveals EV-profiles that are characteristic for the EV donor cell type.

**Figure 3:**
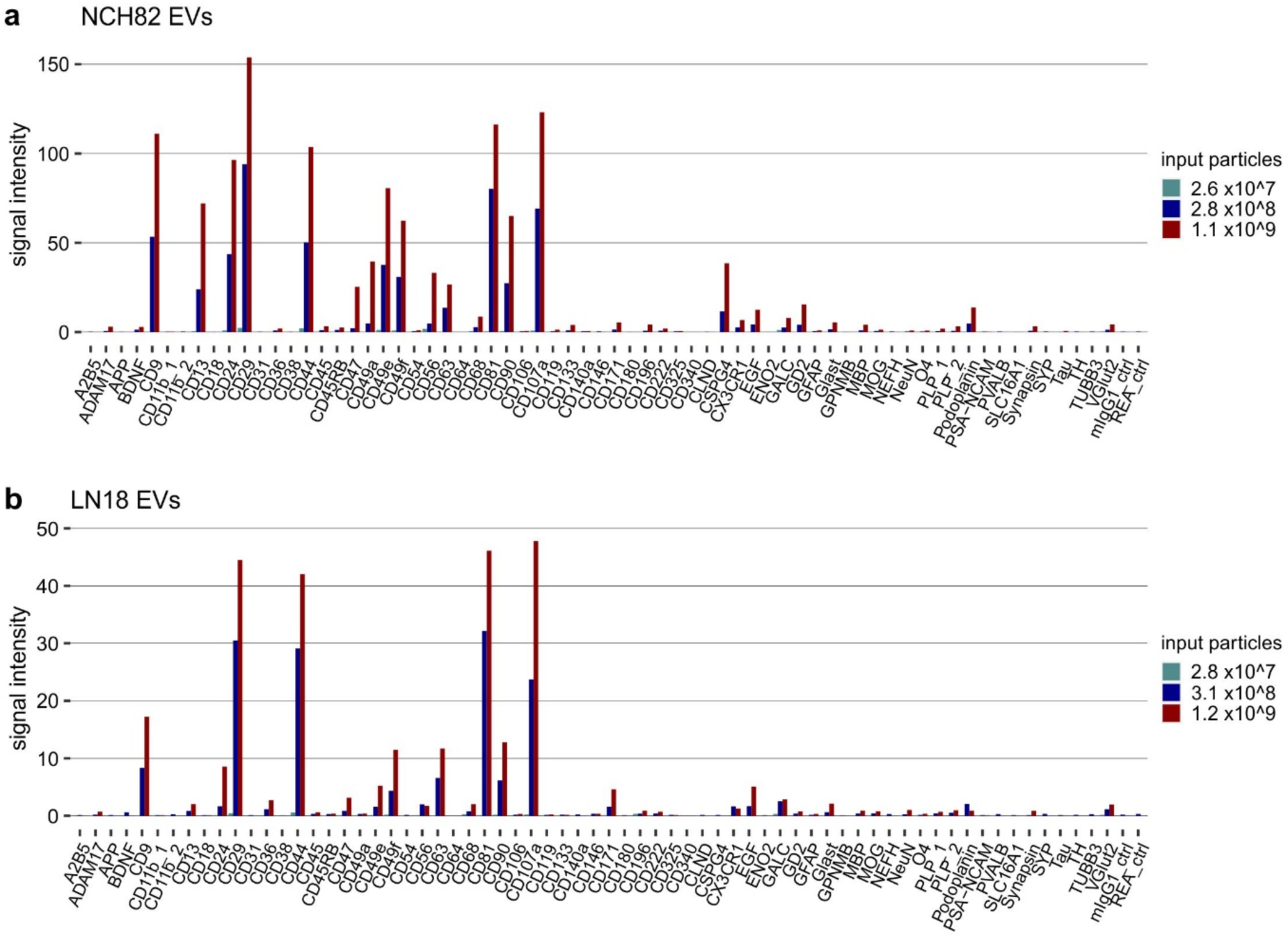
Titration of EVs derived from glioblastoma cells. EV Neuro median signal intensities for increasing total numbers of NCH82 EVs (a) and LN18 EVs (b) as determined by NTA particle count. The number of 2.8×10^7^ particles (turquoise) represents the input of 120 µl of pre-cleared cell culture supernatant (10,000 × g) without prior EV enrichment. Other particle inputs (blue and red) were achieved by EV enrichment via dUC. Bars reflect marker profiles of one experiment each.

### Profiling of GB serum EVs compared to healthy controls

Having verified that the EV Neuro assay depicts differential marker profiles in GB cell-derived EVs and primary human astrocyte-derived EVs, we examined whether similar differences can also be detected between circulating EVs of GB patients (n=9) and healthy controls (n=12). Therefore, we separated EVs from 500 µl serum using SEC and further enrichment by UC (SEC-UC; Fig. S4a, b) followed by EV Neuro analysis. Absolute signal intensities were strongly varying between the individuals tested in both the GB-patient and HC group, although not entirely comparable since underlying serum input volumes per EV Neuro panel slightly differed between subjects (200 ± 50 µl; Fig. S4c). We therefore focused on comparing the EV Neuro marker profiles (normalized signals) of patient- and control-EV samples (Fig. 4a). We observed an increase in the proportion of CD63^+^EVs and CD81^+^EVs in GB patients, while CD9^+^EVs (largely representing platelet EVs in the circulation) remained constant. This relative increase of CD63^+^CD81^+^EVs in GB patients may indicate higher levels of EVs from origins other than platelets.

**Figure 4:**
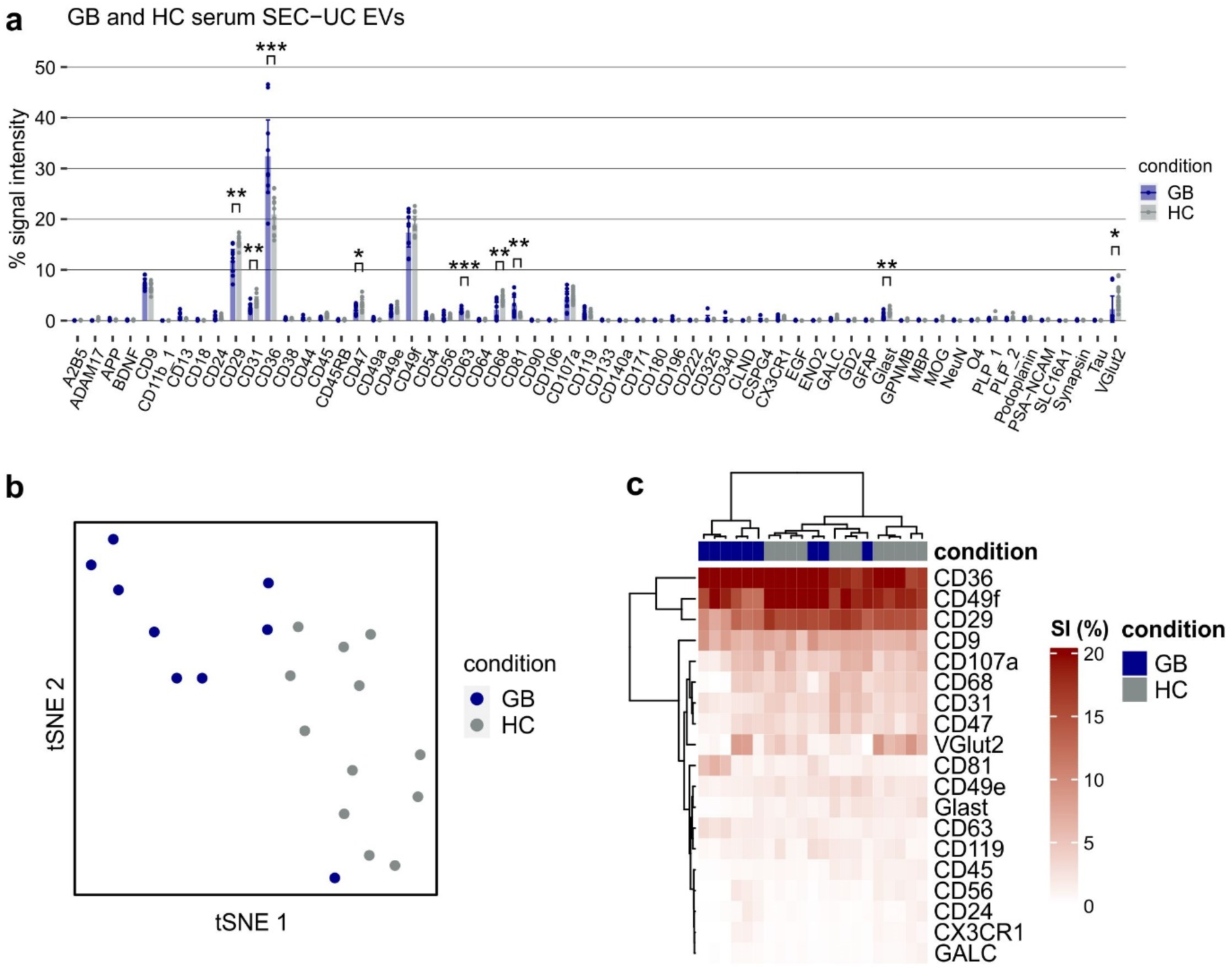
Profiling of serum EVs from GB patients and healthy controls. (a) Relative EV Neuro signal intensities for EVs separated from the serum of GB patients or HC by SEC followed by UC calculated as signal of target divided by the total signal of all markers (in %). Bars represent mean values and error bars indicate the 95 % confidence interval. Asterisks mark statistically significant differences between GB and HC. Only significant alterations of targets that were consistently detected above background in all individuals are marked. *=p<.05, **=p<.01, ***=p<.001. (b) tSNE on data from (a) stratified by condition (color). (c) Heatmap visualization of selected targets from (a) including hierarchical clustering for targets as well as subjects.

Intriguingly, we found a significant increase in the proportion of CD36^+^EVs, which we already found enriched in GB cell-derived EVs (LN229-EVs) compared to astrocytes. None of the other markers that appeared enriched in GB cell line derived EVs, was reappearing in the patients’ samples. Also, no unique marker appeared in the profile of GB patients compared to HC-EV samples. However, the relative intensities of some markers (CD29, CD31, CD47, CD68, GLAST VGlut2) were slightly decreased in GB patients compared to HC, indicating a proportional shift of EVs carrying these markers to the CD36 population. Performing t-distributed stochastic neighbor embedding (tSNE) analysis on the obtained data allowed separation of GB samples from HC samples (Fig. 4b), underscoring that EV profiles in GB and HC are distinct. Heatmap clustering analysis focusing on markers consistently detected above background in all individuals confirms the separation of GB and HC subjects except for three GB patients which cluster within the HC group (Fig. 4c). Taken together, the EV Neuro assay allowed to distinguish GB and HC serum derived EV profiles and specified serum EV associated CD36 as a potential candidate GB biomarker.

### Profiling of MS plasma EVs compared to healthy controls

Next, we tested whether the EV Neuro assay can identify differences between circulating EVs of MS patients and healthy persons and thus be potentially useful for MS biomarker discovery. Here, we used immuno-affinity capture for CD63 or CD81 and magnetic isolation to separate and enrich CD63^+^EVs and CD81^+^EVs, respectively, from each 500 µl of plasma of MS patients (n=9) and HC (n=5). EV Neuro signal intensity analysis for CD63^+^EVs and CD81^+^EVs revealed a large signal variation in MS patient and control EV samples for all analyzed markers despite constant underlying plasma sample input per EV Neuro panel (250 µl), indicating high interindividual variation independent of disease state ([upper graphs in Fig. 5 a (CD63^+^EVs) and b (CD81^+^EVs)]. As a result, a high effect size for a specific target and/or a large sample set is needed to detect differences among absolute signal intensities. Notably, specific and differential analysis of CD63^+^ and CD81^+^EV subpopulations using the EV Neuro assay is technically feasible and produces robust signal intensities.

**Figure 5:**
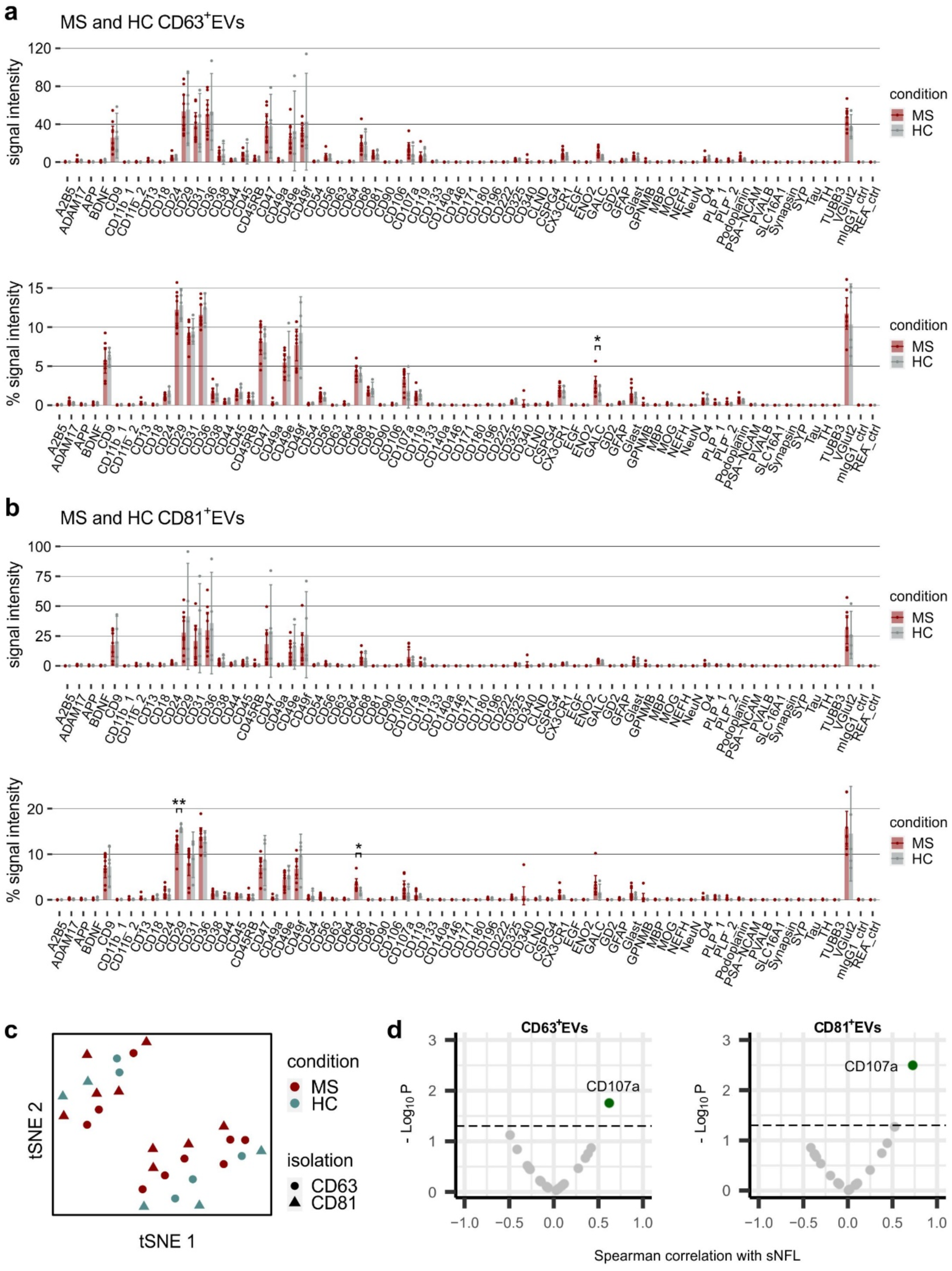
Profiling of plasma EVs from MS patients and healthy controls. EV Neuro signal intensities for EVs separated from the plasma of MS patients or HC by CD63- (a) or CD81- (b) immuno-affinity capture, as well as relative signal intensities as signal of target divided by the total signal of all markers (in %). Bars represent mean values and error bars indicate the 95 % confidence intervals. Asterisks mark statistically significant differences between MS and HC in relative profiles. Only significant alterations of targets that were consistently detected above background in all individuals are marked. *=p<.05. (c) tSNE was performed on relative signal intensities of anti-CD63 and anti-CD81 immuno-affinity captured EVs and stratified by condition (color) and isolation procedure (shape). (d) Volcano blots of Spearman correlation analysis of neurofilament (sNFL) values in blood with relative signal intensities of selected EV Neuro markers in CD63^+^EVs and CD81^+^EVs, respectively (dashed line p =.05).

The illustration of normalized signal intensities shows more uniform EV Neuro profiles compared to absolute signals [lower graphs in Fig. 5 a (CD63^+^EVs) and b (CD81^+^EVs)]. This highlights again that the profile of circulating EVs is comparable in all analyzed persons regardless of individual differences in EV concentration and efficiency of EV isolation. Notably, when comparing the EV profile of MS patients and HC, a few markers revealed significant differences: Galactosylceramide (GALC), a myelin-specific lipid, was increased in CD63^+^EVs of MS patients; CD68 (LAMP4), which is specific for monocytes, macrophages, and microglia, was increased in CD81^+^EVs of MS patients; and CD29 (integrin beta 1) was decreased in CD81^+^EVs of MS patients. Of note, CD107a (LAMP1) showed a clear trend of signal increase in both CD63^+^EVs and CD81^+^EVs. However, performing tSNE data analysis of CD63^+^EVs and CD81^+^EVs in MS and HC did neither result in separation of health condition nor of isolation procedure (Fig. 5c). Thus, general EV profiles do not discriminate the CD63^+^ and CD81^+^EV subpopulations and appear similar in MS and HC, at least in a cross-sectional analysis of small cohorts.

The level of free neurofilament light chain in serum (sNFL) has recently been introduced as biomarker of prognosis and treatment response that may be combined with other liquid biopsy markers (32–35). Therefore, we performed a correlation analysis of sNFL levels with those EV markers consistently detected by the Neuro EV assay and observed a significant correlation of sNFL with EVs positive for CD107a (CD63^+^EVs: rho=0.62, p=0.020, n=14; CD81^+^EVs: rho=0.73, p=0.004, n=14; Fig. 5d). In conclusion, the EV Neuro assay identified potentially disease-relevant EV markers that can be correlated with established EV-independent markers and further validated through follow-up studies.

In the pilot cohort of MS patients used in this study, six patients were in a stable disease phase and three in a disease relapse. Analyzing the marker profiles (relative signal intensities) of CD63^+^EVs and CD81^+^EVs revealed a comparable variation of individual markers (Fig. 6). However, CD49e (integrin alpha 5) in CD63^+^EVs and CD29 (integrin beta 1) in CD81^+^EVs were significantly increased in a disease relapse versus a stable disease phase, though a larger sample size is required for retrieving a more solid statistical evaluation of these markers.

**Figure 6:**
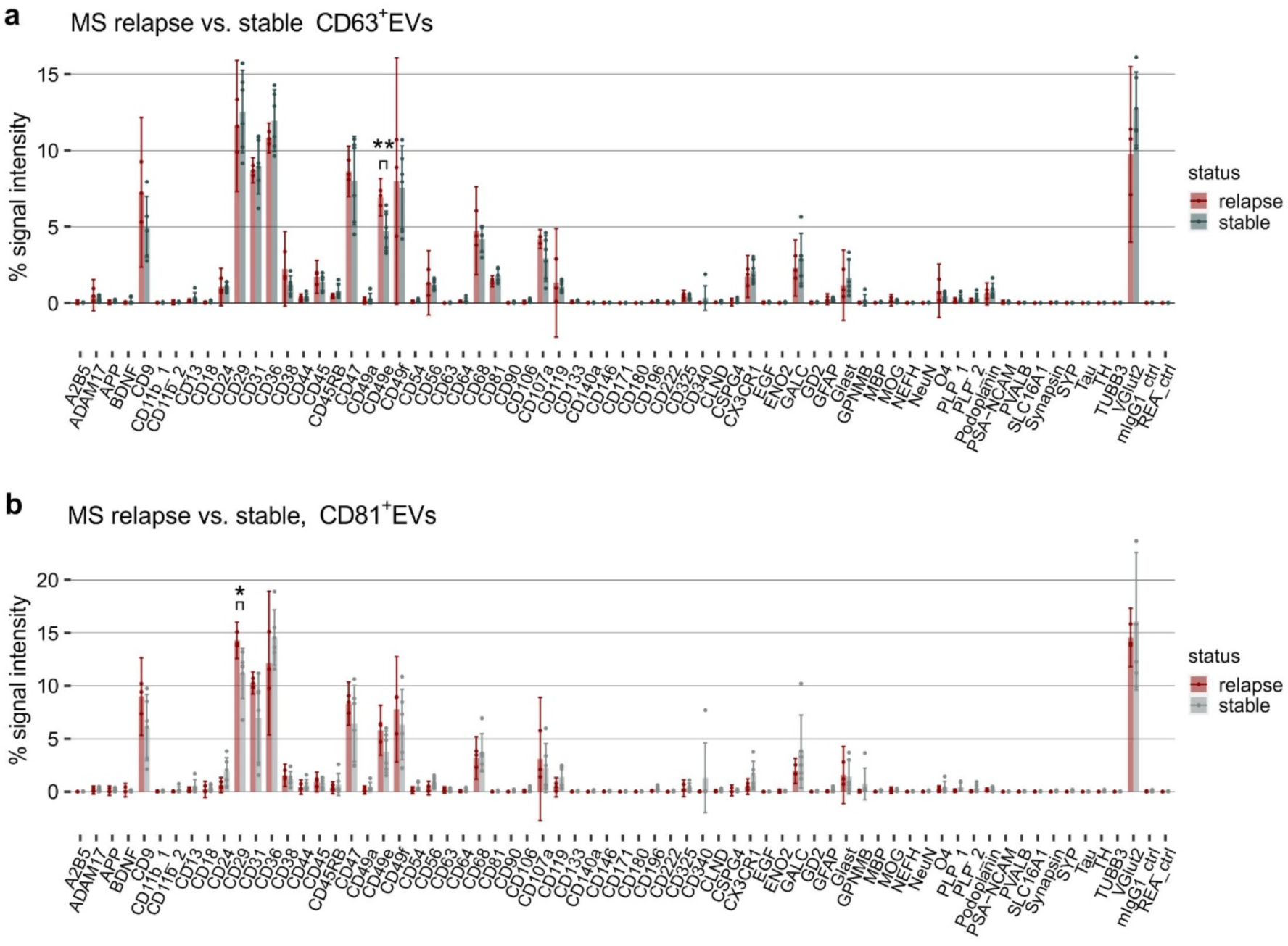
Comparison of plasma EVs from MS patients in a relapse versus in a stable disease phase. Relative EV Neuro signal intensities as signal of target divided by the total signal of all markers (in %) for immuno-affinity captured CD63^+^EVs (a) and CD81^+^EVs (b). Bars represent mean values and error bars indicate the 95 % confidence interval. Asterisks mark statistically significant differences between stable phase and disease relapse. Only significant alterations of targets that were consistently detected above background in all individuals are marked. *=p<.05, **=p<.01.

Taken together, utilizing immuno-capture and subsequent EV Neuro analysis to phenotype EVs in as little as 250 µl of plasma of MS patients and HC enabled EV profiling and identification of candidate biomarkers for MS liquid biopsy.

### Profiling of MCI/AD/DEP plasma-derived EVs

To test further marker reactivity of the EV Neuro assay beyond GB and MS pathology, we included a cohort of patients (MAD, n=8) suffering from mild cognitive impairment, Alzheimer’s disease, depression, or a combination of the named. We used immuno-affinity capture for CD81 and magnetic isolation to separate and enrich CD81^+^EVs from 1 ml plasma of the patients (corresponds to 500 µl per EV Neuro panel). As observed for other cohorts, a high interindividual variation in absolute signal intensities was observed (Fig. S5), but the relative profiles of the patients were more uniform (Fig.7a). The same markers as in the MS/HC cohort turned up, while markers specific for the MAD condition that could be of potential biomarker interest were not detected within the small cohort. MAD samples and MS/HC plasma samples were obtained in different laboratories, which could introduce a preanalytical bias (e.g., grade of platelet contamination). Nevertheless, to compare the EV profiles of the MAD condition with the other conditions evaluated in this study, we performed tSNE analysis (Fig. 7b). Interestingly, assessment of MAD, MS, and HC relative signals showed a clear separation of the MAD cohort, which was confirmed by heatmap clustering analysis of markers consistently detected above background (Fig. 7c). However, comprehensive heatmap cluster analysis of normalized EV Neuro data including all blood-derived EV samples (including GB serum) and markers analyzed showed separation of the samples predominantly by the blood collection type, isolation method, and the lab of blood collection, while the health condition was a comparably less stable parameter (Fig. S6). These findings underscore that using the Neuro EV assay for reliable biomarker identification requires a strict control of preanalytical parameters, as generally recommended for the analysis of blood EVs (36–38).

**Figure 7:**
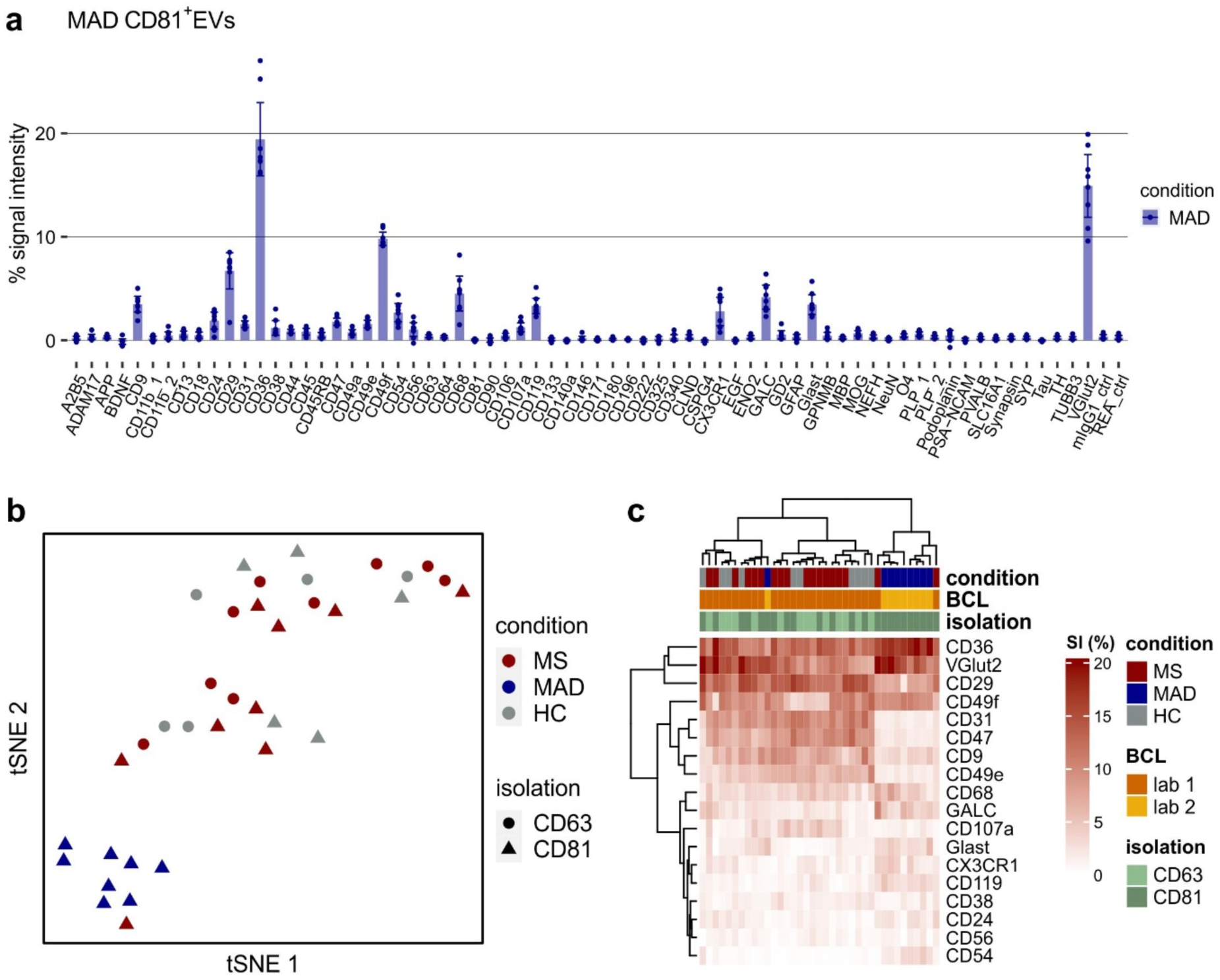
Phenotyping of plasma EVs from MAD patients and comparison of all plasma samples. (a) Relative EV Neuro signal intensities for EVs separated from the plasma of MAD patients by immuno-affinity capture (anti-CD81) and magnetic separation. Bars represent mean values and error bars indicate the 95 % confidence intervals (n=8 patients). (b) tSNE of relative signal intensities from all plasma-derived EV samples analyzed (MS, MAD, HC) stratified by condition (color) and isolation (shape). (c) Heatmap visualization of relative signals for selected targets from all plasma samples analyzed including hierarchical clustering for targets as well as subjects (BCL = lab of blood collection).

## Discussion

In this study, we assessed the performance of a novel multiplex bead-based flow cytometry platform (a prototype of the MACSPlex EV Kit Neuro) for detecting brain-derived EVs using cell culture supernatant, serum, and plasma as starting materials and different methods of EV isolation. The platform was able to detect differences in EV numbers and to reveal changes of EV-populations within an EV profile, which enables its use as a starting point to detect potential biomarkers among a pre-selection of EV-associated markers. In small pilot cohorts of patient-derived blood samples, including GB, MS, and AD, it was possible to perform EV profiling and to reveal potential biomarker candidates that could be followed up in larger cohorts and by using other detection technologies.

Using EVs collected from GB cell line and human primary astrocyte cell culture-derived EVs, we performed a qualitative validation of individual EV-markers within the EV Neuro assay and a basic evaluation of their quantitative range. Marker detection was dependent on the EV input and signals appeared semi-quantitative with non-linear increases, in particular at higher signal intensities. Signal saturation is expected with such a bead-based assay, where the number of binding sites on the beads and steric capacity is limited. Plasma and serum samples produced a recurrent EV-profile with high interindividual variation in signal intensities of the markers, illustrating that high effect sizes and large sample cohorts are required to obtain reliable statistical outcomes in cross-sectional biomarker studies. Longitudinal studies of patient samples, though, may be promising to reveal changes within an individual’s EV marker signal intensities, which indeed has been achieved previously with the same technology using another marker portfolio representing circulation markers (24). Notably, the EV-markers detected in blood and especially those revealing the dominant signals were not CNS-specific. Markers like L1CAM/CD171, A2B5, PLP, MOG, which are expected to be present on the surface of neuronal or glial EVs (39,40), were never detected with reliable sensitivity. Lacking signals of these markers could be either due to the absence of these EVs in the sample, their presence below the limit of detection, or poor performance of the bead-linked antibodies (technical limitation). Intracellular markers in the test panel (e.g., NeuN, GFAP, MBP) were never detected, likely due to topology. Therefore, it remains unclear whether the EV Neuro assay can pick up EVs originating from the CNS in the circulation. However, the detection of EV profiles in the circulation that deviate from healthy controls and correlate with CNS disease appears feasible. The sample input required per EV Neuro panel was minimal with 150 to 500 µl of plasma or serum and within the clinically practical range.

To deal with high interindividual variation in absolute signal intensities, we normalized individual markers over the total of all retrieved signals, expressing their relative representation within the group of depicted markers. This normalization strategy allows comparison of EV-profiles independent of parameters such as sample input or individual variations in total EV counts and was employed previously (25). Identification of markers sticking out of the relative profile may help to recognize EV populations of functional relevance or of biomarker value and to stratify patients according to disease condition. Next to assessing the relative representation of the markers, other normalization strategies can be useful depending on the research question. Normalization on one or all three of the tetraspanins CD9, CD63, and CD81 suggested by the kit’s manual may provide additional information on the composition of the complex EV profiles [e.g., in (26)]. In our study, the relative profiles of EVs released by different GB cell lines were distinct and the assay indicated changes of EV markers in the serum and plasma of GB and MS patients versus controls. Notably, tSNE analysis almost completely separated GB EV-profiles from healthy profiles, indicating a relevant shift in markers within the profile. However, clustering analysis indicated that sampled individuals not only group according to disease condition, but technical factors such as the lab of blood collection and the EV isolation procedure were at least equally relevant for clustering. Preanalytical parameters must therefore be tightly controlled (36–38), control samples should be carefully chosen, and large sample sizes are important for identifying disease-specific changes in individual EV markers within the complex mixture of circulating EVs.

Although the EV Neuro analysis in this study was based on a small number of samples, markers of potentially relevant EV populations could be visualized. For example, CD36^+^EVs were elevated in cell line-derived EVs as well as in serum derived EVs from GB patients. It has been suggested that CD36 regulates glioblastoma cell migration and proliferation, and preliminary data indicate that low CD36 levels in glioblastoma patients may be associated with a better prognosis (41). Moreover, GalC^+^EVs and CD68^+^EVs were found slightly elevated in the plasma of MS patients. Since GalC is an oligodendrocyte/myelin marker and CD68 is a macrophage-lineage marker, these observations may reflect an increased presence of oligodendrocyte-derived EVs and microglia/monocyte-derived EVs in the circulation, respectively. Although these markers seem to be related to the biological background of the disease, the limited sample size does not allow to draw definite conclusions but might provide some guidance for follow-up analysis.

The study represents an exploratory analysis using a range of different sample materials to evaluate the potential of the MACSPlex EV Kit Neuro to be used for detecting CNS-derived EVs in the context of disease focusing on qualitative aspects. Further studies need to be performed to reveal more detailed information regarding the limit of detection, the range of linearity, and the point of saturation (21,22). Furthermore, validation of the marker performance on specified sample material was limited to EVs of astrocytic and GB origin and may be expanded to EVs originating from other human neuronal and glial cells, as well as from brain microvascular endothelial cells. A limitation of the EV Neuro assay per se may be that the marker panel to be included in the assay platform is restricted to 37 different surface markers and requires a pre-selection according to scientific and technical criteria. We used here a beta-test version of the assay that will be refined regarding its marker composition before becoming commercially available. In principle, further markers can be addressed by the assay by varying the detection antibody (21,23).

## Conclusions

In conclusion, the MACSPlex Neuro EV platform is suitable for the assessment of multiparametric EV profiles and the semi-quantitative detection of individual markers in diverse EV sample material including small volumes of body fluids. The assay has demonstrated considerable potential for initial discovery of EV-associated biomarkers and monitoring of circulating EV-profiles related to various CNS disease states. Phenotyping and semi-quantitative profiling of EVs associated with CNS diseases will represent a major step forward for biomarker discovery and clinical research.

## Acknowledgements

We thank all blood donors for their participation and support.

## Data availability

The datasets used and analyzed during the current study are available from the corresponding authors on reasonable request.

## Ethics approval

The experimental procedures were approved by the Human Ethics Committee Rhineland-Palatinate (for MS patients and HC: protocol number 837.019.10) or by the local ethics committee of the University of Bonn (Protocol number for glioblastoma patients Bonn: 182/08; healthy volunteers: 007/17), respectively. The Alzheimer patients were recruited from daily clinical work after having given written consent (§ 14 AVB, University Medical Center Johannes Gutenberg-University Mainz). The experimental procedures adhere to the standards of the Declaration of Helsinki of the World Medical Association.

## Patient consent

Written informed consent to participate was obtained from all patients and healthy persons.

## References

1. Thompson AG, Gray E, Heman-Ackah SM, Mäger I, Talbot K, Andaloussi SE, u. a. Extracellular vesicles in neurodegenerative disease - pathogenesis to biomarkers. Nat Rev Neurol. Juni 2016;12(6):346–57.

2. Vandendriessche C, Bruggeman A, Van Cauwenberghe C, Vandenbroucke RE. Extracellular Vesicles in Alzheimer’s and Parkinson’s Disease: Small Entities with Large Consequences. Cells. 15. November 2020;9(11):2485.

3. Macedo-Pereira A, Martins C, Lima J, Sarmento B. Digging the intercellular crosstalk via extracellular vesicles: May exosomes be the drug delivery solution for target glioblastoma? J Control Release Off J Control Release Soc. Juni 2023;358:98–115.

4. Van Niel G, D’Angelo G, Raposo G. Shedding light on the cell biology of extracellular vesicles. Nat Rev Mol Cell Biol. 2018;19(4):213–28.

5. Saint-Pol J, Gosselet F, Duban-Deweer S, Pottiez G, Karamanos Y. Targeting and Crossing the Blood-Brain Barrier with Extracellular Vesicles. Cells. 1. April 2020;9(4):851.

6. Busatto S, Morad G, Guo P, Moses MA. The role of extracellular vesicles in the physiological and pathological regulation of the blood-brain barrier. FASEB BioAdvances. September 2021;3(9):665–75.

7. Krämer-Albers EM. Extracellular Vesicles at CNS barriers: Mode of action. Curr Opin Neurobiol. August 2022;75:102569.

8. Ostrom QT, Price M, Neff C, Cioffi G, Waite KA, Kruchko C, u. a. CBTRUS Statistical Report: Primary Brain and Other Central Nervous System Tumors Diagnosed in the United States in 2015-2019. Neuro-Oncol. 5. Oktober 2022;24(Suppl 5):v1–95.

9. Tzaridis T, Weller J, Bachurski D, Shakeri F, Schaub C, Hau P, u. a. A novel serum extracellular vesicle protein signature to monitor glioblastoma tumor progression. Int J Cancer. 15. Januar 2023;152(2):308–19.

10. Mycko MP, Baranzini SE. microRNA and exosome profiling in multiple sclerosis. Mult Scler Houndmills Basingstoke Engl. April 2020;26(5):599–604.

11. Aharon A, Spector P, Ahmad RS, Horrany N, Sabbach A, Brenner B, u. a. Extracellular Vesicles of Alzheimer’s Disease Patients as a Biomarker for Disease Progression. Mol Neurobiol. Oktober 2020;57(10):4156–69.

12. Mazzucco M, Mannheim W, Shetty SV, Linden JR. CNS endothelial derived extracellular vesicles are biomarkers of active disease in multiple sclerosis. Fluids Barriers CNS. 8. Februar 2022;19(1):13.

13. Marostica G, Gelibter S, Gironi M, Nigro A, Furlan R. Extracellular Vesicles in Neuroinflammation. Front Cell Dev Biol. 2020;8:623039.

14. Simonsen JB. What are we looking at? Extracellular vesicles, lipoproteins, or both? Circ Res. 2017;

15. Fiandaca MS, Kapogiannis D, Mapstone M, Boxer A, Eitan E, o u. a. Identification of preclinical Alzheimer’s disease by a profile of pathogenic proteins in neurally derived blood exosomes: A case-control study. Alzheimers Dement J Alzheimers Assoc. Juni 2015;11(6):600–607.e1.

16. Jiang C, Hopfner F, Berg D, Hu MT, Pilotto A, Borroni B, u. a. Validation of α-Synuclein in L1CAM-Immunocaptured Exosomes as a Biomarker for the Stratification of Parkinsonian Syndromes. Mov Disord Off J Mov Disord Soc. November 2021;36(11):2663–9.

17. Pulliam L, Sun B, Mustapic M, Chawla S, Kapogiannis D. Plasma neuronal exosomes serve as biomarkers of cognitive impairment in HIV infection and Alzheimer’s disease. J Neurovirol. Oktober 2019;25(5):702–9.

18. Goetzl EJ, Abner EL, Jicha GA, Kapogiannis D, Schwartz JB. Declining levels of functionally specialized synaptic proteins in plasma neuronal exosomes with progression of Alzheimer’s disease. FASEB J. 2018;32(2):888–93.

19. Norman M, Ter-Ovanesyan D, Trieu W, Lazarovits R, Kowal EJK, Lee JH, u. a. L1CAM is not associated with extracellular vesicles in human cerebrospinal fluid or plasma. Nat Methods. Juni 2021;18(6):631–4.

20. Gomes DE, Witwer KW. L1CAM-associated extracellular vesicles: A systematic review of nomenclature, sources, separation, and characterization. J Extracell Biol. März 2022;1(3):e35.

21. Koliha N, Wiencek Y, Heider U, Jüngst C, Kladt N, Krauthäuser S, u. a. A novel multiplex bead-based platform highlights the diversity of extracellular vesicles. J Extracell Vesicles. 2016;5(1).

22. Wiklander OPB, Bostancioglu RB, Welsh JA, Zickler AM, Murke F, Corso G, u. a. Systematic methodological evaluation of a multiplex bead-based flow cytometry assay for detection of extracellular vesicle surface signatures. Front Immunol. 13. Juni 2018;9(JUN).

23. Welsh JA, Killingsworth B, Kepley J, Traynor T, Cook S, Savage J, u. a. MPAPASS software enables stitched multiplex, multidimensional EV repertoire analysis and a standard framework for reporting bead-based assays. Cell Rep Methods. 24. Januar 2022;2(1):100136.

24. Brahmer A, Neuberger E, Esch-Heisser L, Haller N, Jorgensen MM, Baek R, u. a. Platelets, endothelial cells and leukocytes contribute to the exercise-triggered release of extracellular vesicles into the circulation. J Extracell Vesicles [Internet]. 2019;8(1). Verfügbar unter: https://doi.org/10.1080/20013078.2019.1615820

25. Giovanazzi A, van Herwijnen MJC, Kleinjan M, van der Meulen GN, Wauben MHM. Surface protein profiling of milk and serum extracellular vesicles unveils body fluid-specific signatures. Sci Rep. 30. Mai 2023;13(1):8758.

26. Ekström K, Crescitelli R, Pétursson HI, Johansson J, Lässer C, Olofsson Bagge R. Characterization of surface markers on extracellular vesicles isolated from lymphatic exudate from patients with breast cancer. BMC Cancer. 10. Januar 2022;22(1):50.

27. Van Deun J, Mestdagh P, Agostinis P, Akay Ö, Anand S, Anckaert J, u. a. EV-TRACK: Transparent reporting and centralizing knowledge in extracellular vesicle research. Nat Methods. 2017;14(3):228–32.

28. Schiffer D, Mellai M, Boldorini R, Bisogno I, Grifoni S, Corona C, u. a. The Significance of Chondroitin Sulfate Proteoglycan 4 (CSPG4) in Human Gliomas. Int J Mol Sci. 12. September 2018;19(9):2724.

29. Nazha B, Inal C, Owonikoko TK. Disialoganglioside GD2 Expression in Solid Tumors and Role as a Target for Cancer Therapy. Front Oncol. 2020;10:1000.

30. Chen C, Zhao S, Karnad A, Freeman JW. The biology and role of CD44 in cancer progression: therapeutic implications. J Hematol OncolJ Hematol Oncol. 10. Mai 2018;11(1):64.

31. Wang J, Li Y. CD36 tango in cancer: signaling pathways and functions. Theranostics. 2019;9(17):4893–908.

32. Bittner S, Steffen F, Uphaus T, Muthuraman M, Fleischer V, Salmen A, u. a. Clinical implications of serum neurofilament in newly diagnosed MS patients: A longitudinal multicentre cohort study. EBioMedicine. Juni 2020;56:102807.

33. Bittner S, Oh J, Havrdová EK, Tintoré M, Zipp F. The potential of serum neurofilament as biomarker for multiple sclerosis. Brain J Neurol. 29. November 2021;144(10):2954–63.

34. Benkert P, Meier S, Schaedelin S, Manouchehrinia A, Yaldizli Ö, Maceski A, u. a. Serum neurofilament light chain for individual prognostication of disease activity in people with multiple sclerosis: a retrospective modelling and validation study. Lancet Neurol. März 2022;21(3):246–57.

35. Thebault S, Bose G, Booth R, Freedman MS. Serum neurofilament light in MS: The first true blood-based biomarker? Mult Scler Houndmills Basingstoke Engl. September 2022;28(10):1491– 7.

36. Coumans FAW, Brisson AR, Buzas EI, Dignat-George F, Drees EEE, El-Andaloussi S, u. a. Methodological Guidelines to Study Extracellular Vesicles. Circ Res. 12. Mai 2017;120(10):1632– 48.

37. Clayton A, Boilard E, Buzas EI, Cheng L, Falcón-Perez JM, Gardiner C, u. a. Considerations towards a roadmap for collection, handling and storage of blood extracellular vesicles. J Extracell Vesicles. 2019;8(1):1647027.

38. Théry C, Witwer KW, Aikawa E, Alcaraz MJ, Anderson JD, Andriantsitohaina R, u. a. Minimal information for studies of extracellular vesicles 2018 (MISEV2018): a position statement of the International Society for Extracellular Vesicles and update of the MISEV2014 guidelines. J Extracell Vesicles. 2018;7(1).

39. Fauré J, Lachenal G, Court M, Hirrlinger J, Chatellard-Causse C, Blot B, u. a. Exosomes are released by cultured cortical neurones. Mol Cell Neurosci. 2006;31(4):642–8.

40. Krämer-Albers EM, Bretz N, Tenzer S, Winterstein C, Möbius W, Berger H, u. a. Oligodendrocytes secrete exosomes containing major myelin and stress-protective proteins: Trophic support for axons? Proteomics - Clin Appl. 2007;1(11):1446–61.

41. Hale JS, Otvos B, Sinyuk M, Alvarado AG, Hitomi M, Stoltz K, u. a. Cancer stem cell-specific scavenger receptor CD36 drives glioblastoma progression. Stem Cells Dayt Ohio. Juli 2014;32(7):1746–58.

