## Supplemental Material for "Assessment of technical and clinical utility of a bead-based flow cytometry platform for multiparametric phenotyping of CNS-derived extracellular vesicles"

**Table S1: Capture bead composition of the MACSPlex EV Kit Neuro, human, panel A and B.**

| panel | target | | panel | target |
|---|---|---|---|---|
| A | CD68 | | B | CD18 |
| A | CD340 | | B | CD64 |
| A | CD49a | | B | CD107a |
| A | CD140a | | B | CD180 |
| A | CD171 | | B | CD196 |
| A | CD56 | | B | ADAM17 |
| A | CD13 | | B | APP |
| A | BDNF | | B | CLND |
| A | CD31 | | B | ENO2 |
| A | GFAP | | B | GALC |
| A | CD222 | | B | GPNMB |
| A | CD11b_2 | | B | MOG |
| A | CD24 | | B | PLP_1 |
| A | CD45 | | B | SLC16A1 |
| A | CD133 | | B | NeuN |
| A | O4 | | B | Synapsin |
| A | Glast | | B | Tau |
| A | CD90 | | B | PLP_2 |
| A | CX3CR1 | | B | MBP |
| A | EGF | | B | VGlut2 |
| A | CD325 | | B | TH |
| A | Podoplanin | | B | REA_ctrl |
| A | A2B5 | | B | GD2 |
| A | CD29 | | B | mIgG1_ctrl |
| A | CSPG4 | | B | CD146 |
| A | CD9 | | B | CD11b_1 |
| A | CD63 | | B | NEFH |
| A | CD81 | | B | PVALB |
| A | CD47 | | B | TUBB3 |
| A | PSA-NCAM | | B | SYP |
| A | CD44 | | | |
| A | CD38 | | | |
| A | CD49f | | | |
| A | CD106 | | | |
| A | CD119 | | | |
| A | CD49e | | | |
| A | CD54 | | | |
| A | CD36 | | | |
| A | CD45RB | | | |

**Figure S1**

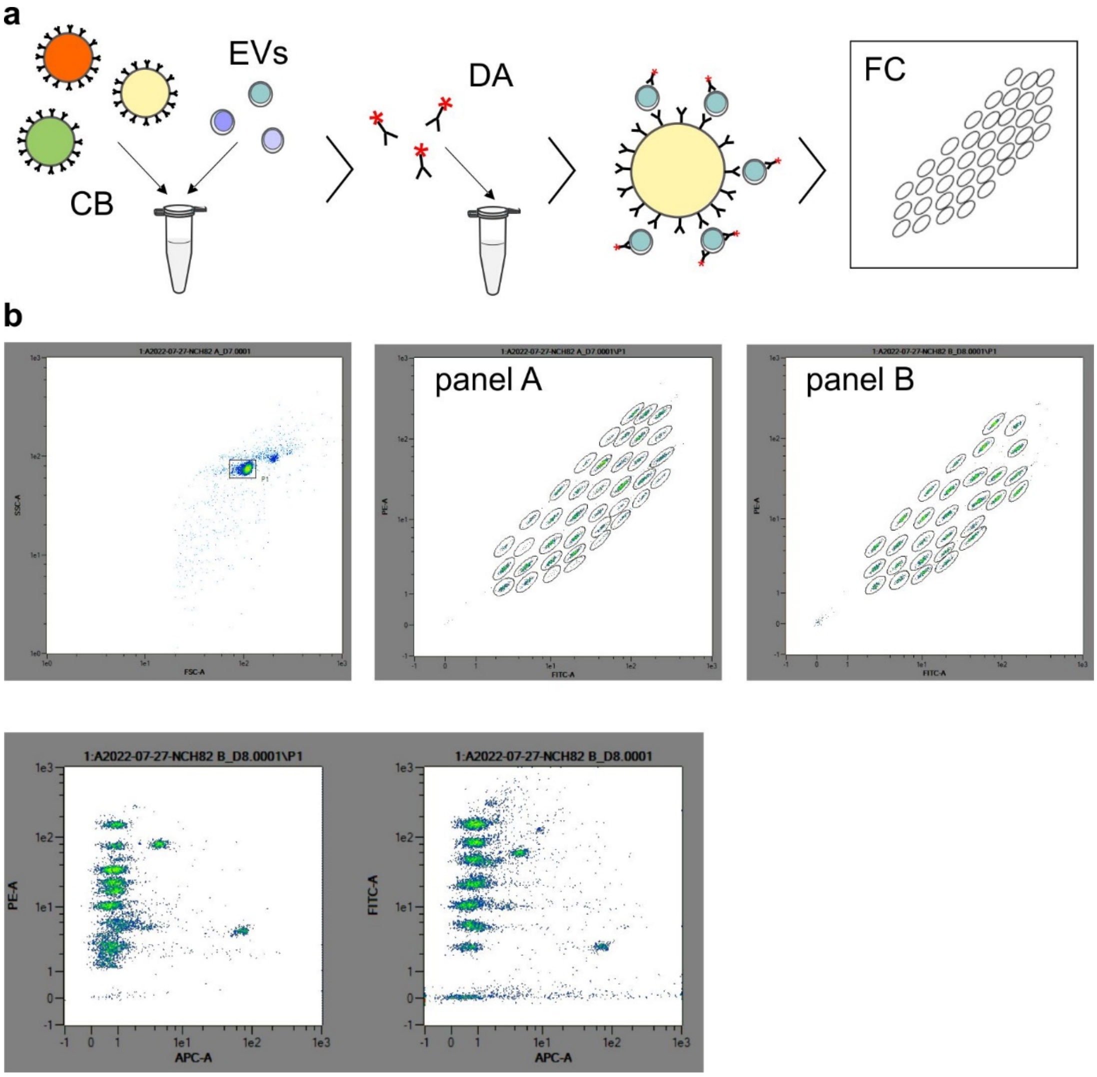


**Figure S1: Workflow and gating strategy for flow cytometry analysis of EV Neuro panel A and B.** (a) Capture beads (CB) are incubated with isolated EVs. After washing, detection antibody cocktail (DA) containing APC-labelled antibodies against CD9, CD63, and CD81 are added. After incubation and washing, flow cytometry (FC) measurement is performed. (b) Gating strategy for flow cytometry assessment of signal intensities for each capture bead population in EV Neuro panel A and B.

**Figure S2**

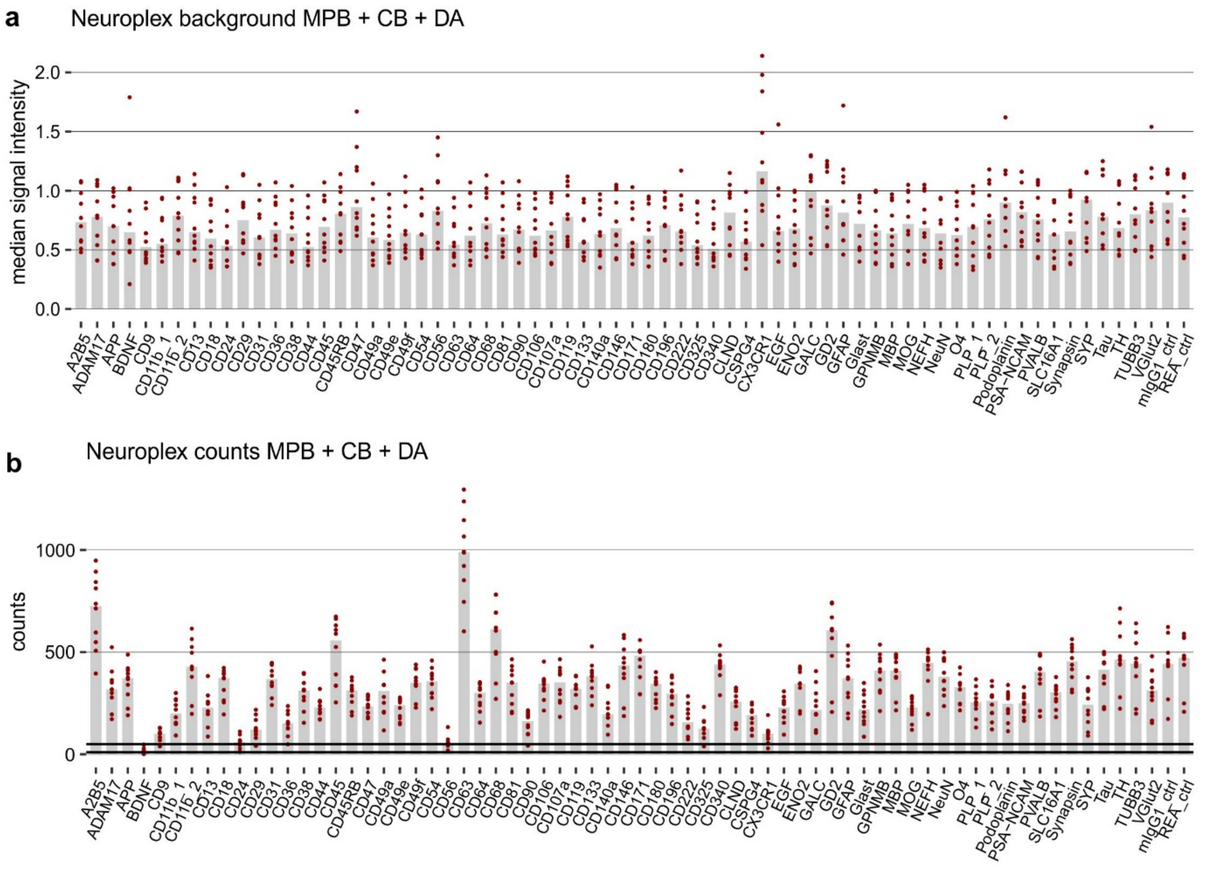


**Figure S2: Number of detected events and background signals.** (a) Background median signal intensities for each capture bead population (CB) after incubation with detection antibody cocktail (DA). Data was obtained from 10 individual measurements of MACSPlex buffer (MPB) with CB and anti-CD9/anti-CD63/anti-CD81-APC DA. Medians are represented as bars. (b) Number of detected barcoded beads used for estimation of median signal intensities (upper bold horizontal line: 50 events, lower bold horizontal line: 10 events). Note: While for most of the targets the number of detected events was consistently between 200 and 500, for BDNF, CD24, and CD56 it was repeatedly below 50 events (for BDNF several times below 10). This results in a small number of events considered to estimate the median signal intensities, which might hamper reproducibility of obtained data for these targets.

**Figure S3**

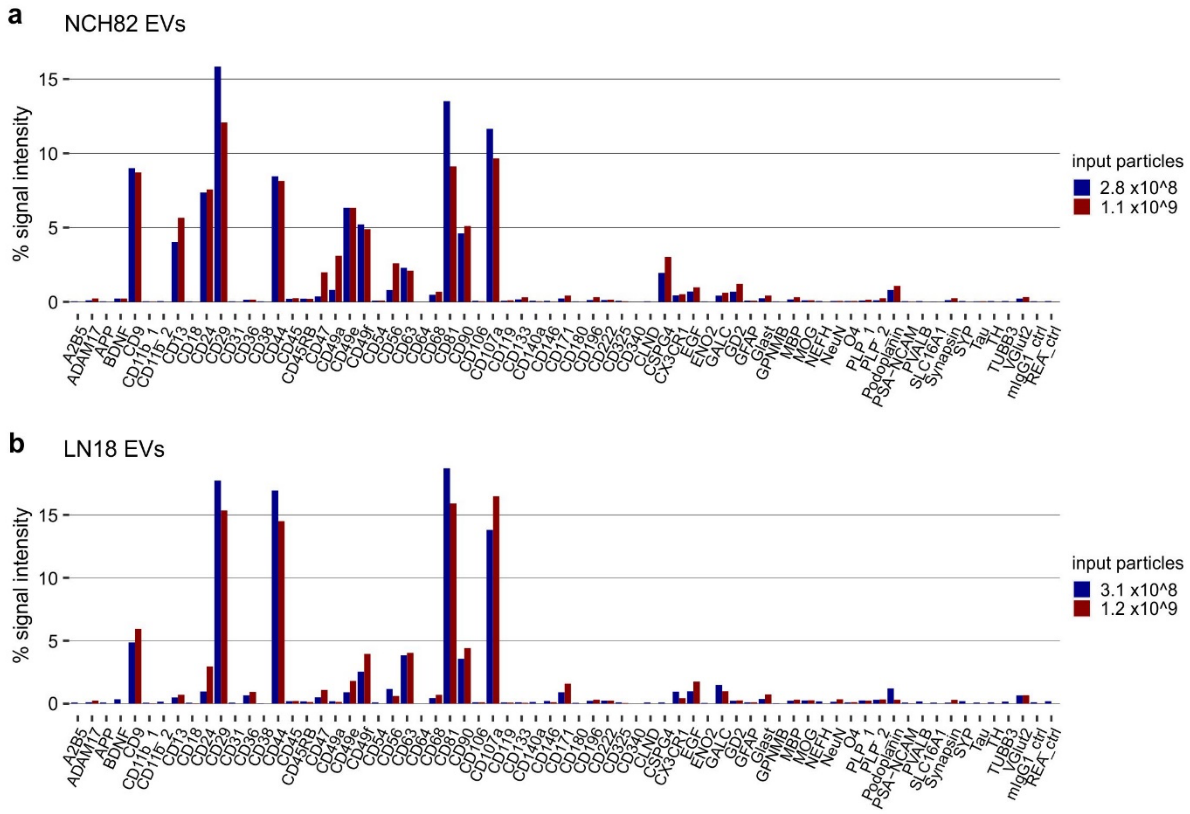


**Figure S3: Titration of EVs derived from glioblastoma cells – relative profiles.** Relative signal intensities achieved with different inputs of NCH82 EVs (a) and LN18 EVs (b) corresponding to Fig. 3 (a) and (b), respectively. The graphs illustrate that despite different EV input, the EV profile remains stable for a specific cell line.

### Figure S4

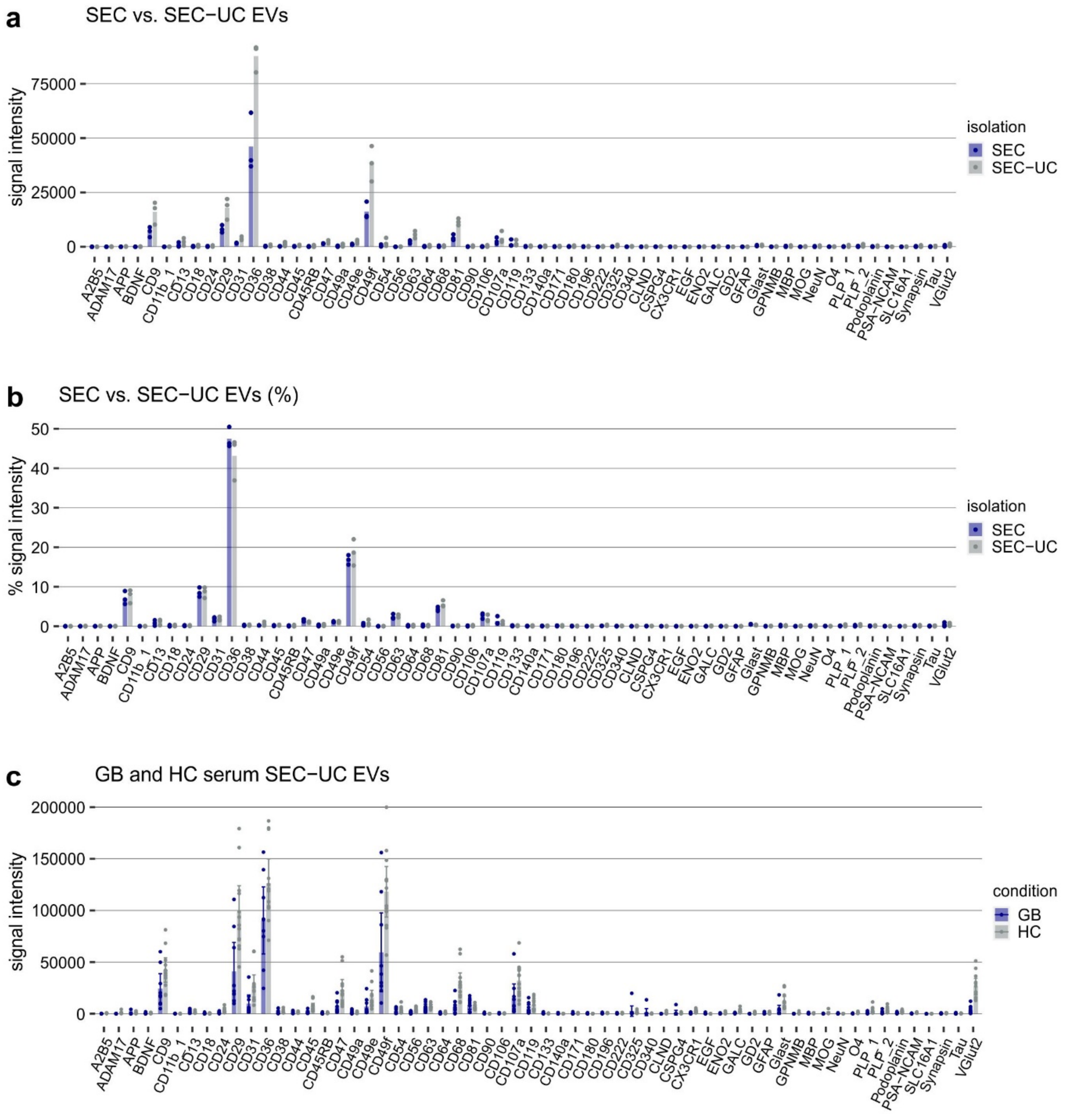


**Figure S4: Phenotyping of serum EVs from GB patients and healthy controls.** (a) EV Neuro signal intensities for EVs separated from the serum of three GB patients by either SEC or combination of SEC and UC (SEC-UC). (b) Relative signal intensities as signal of target divided by the total signal of all markers (in %) of (a). Bars represent mean values. The graphs show that appending UC to SEC leads to increased signal intensity with a constant EV profile. (c) EV Neuro signal intensities for EVs separated from the serum of GB patients or healthy controls by SEC-UC. Corresponding to Fig. 4 in the main text. Note that different input volumes were tested in the GB and HC group, thus the means are not comparable. The graph illustrates the high variation of signal intensities between the individuals of a group and the absence of specific markers unique for the GB condition (or vice versa).

### Figure S5

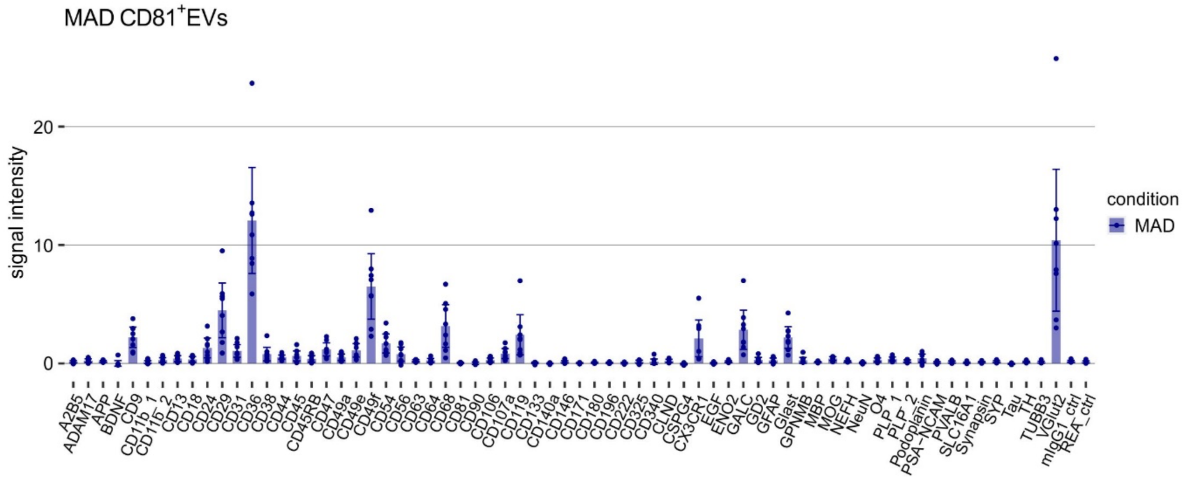


**Figure S5:** EV Neuro signal intensities for EVs separated from the serum of MAD patients by CD81-immunoaffinity capture. Corresponding to Fig. 7a in the main text.

**Figure S6**

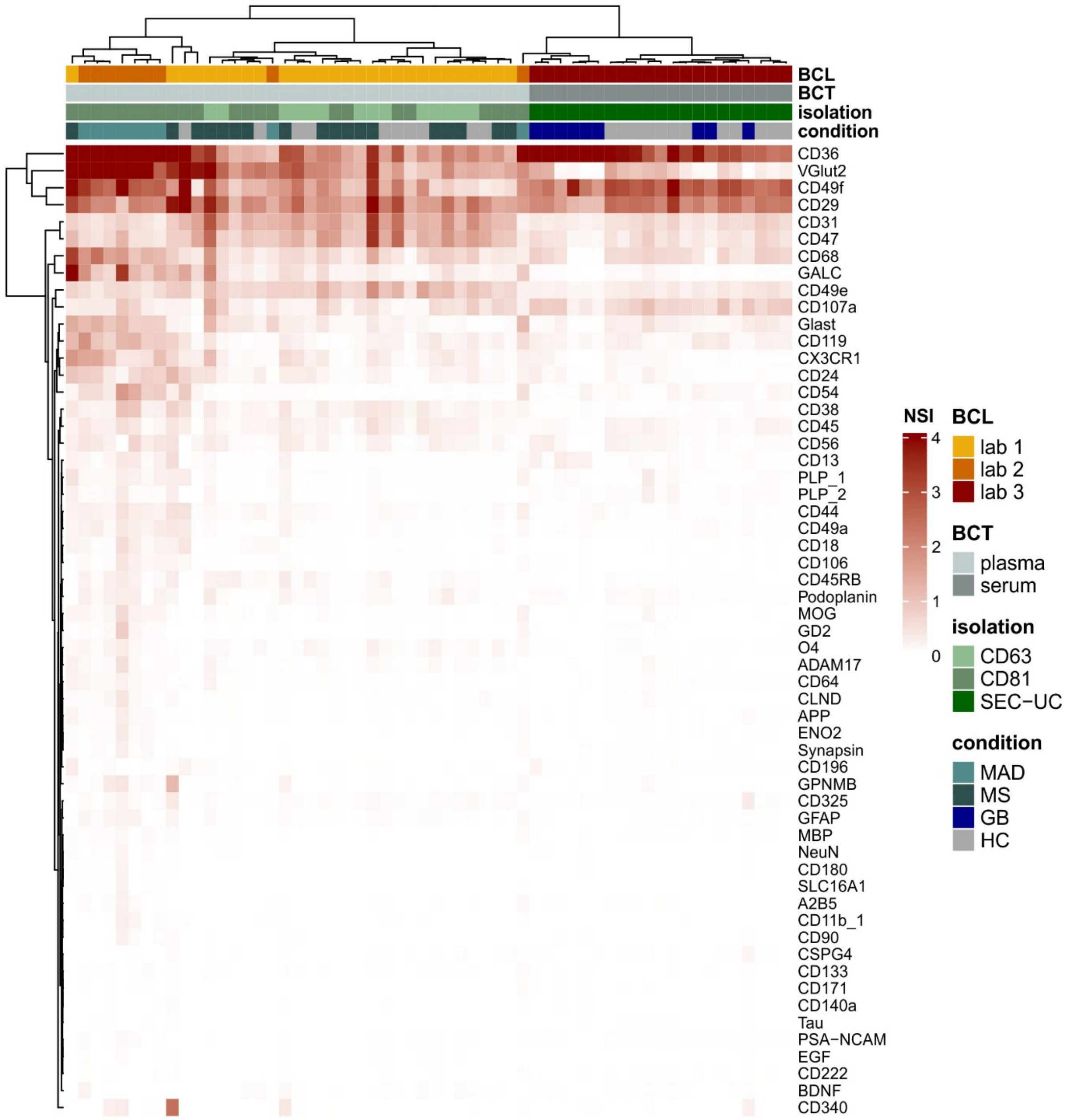


**Figure S6: Heatmap clustering analysis of all blood-derived EV samples.** (a) Heatmap and clustering analysis of all plasma and serum samples analyzed in this study. Signal intensities were normalized to the signal of CD9 (NSI). The genuine EV marker CD9 was used for normalization since it was not affected by the isolation technique and appeared quite stable throughout all samples analyzed. BCL = lab of blood collection, BCT = type of blood collection.
